## Supplementary material for "Transcriptional and genomic parallels between the monoxenous parasite *Herpetomonas muscarum* and *Leishmania*": Fig S1

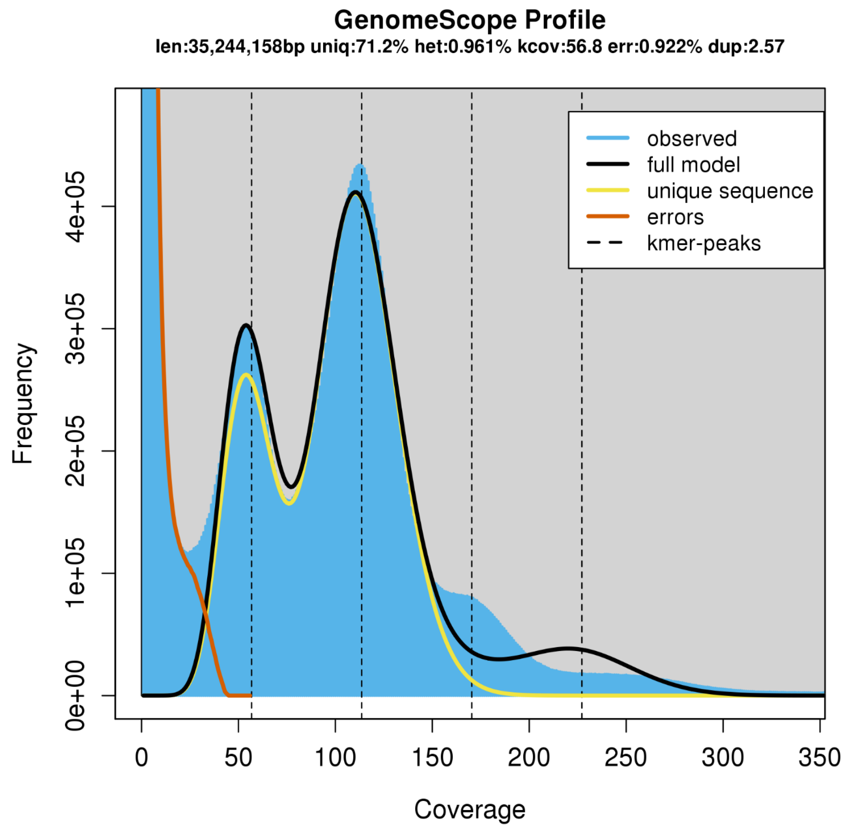


k = 21

```
property min max
Heterozygosity 0.943684% 0.978345%
Genome Haploid Length 35,065,026 bp 35,244,158 bp
Genome Repeat Length 10,090,824 bp 10,142,374 bp
Genome Unique Length 24,974,201 bp 25,101,784 bp
Model Fit 87.3153% 88.9358%
Read Error Rate 0.921767% 0.921767%


Formula: y ~ (((2 * (1 - d) * (1 - (1 - r)^k)) + (2 * d * (1 - (1 - r)^k)^2) +
    (2 * d * ((1 - r)^k) * (1 - (1 - r)^k))) * dnbinom(x, size = kmercov/bias,
    mu = kmercov) * length + (((1 - d) * ((1 - r)^k)) + (d *
    (1 - (1 - r)^k)^2)) * dnbinom(x, size = kmercov * 2/bias,
    mu = kmercov * 2) * length + (2 * d * ((1 - r)^k) * (1 -
    (1 - r)^k)) * dnbinom(x, size = kmercov * 3/bias, mu = kmercov *
    3) * length + (d * (1 - r)^(2 * k)) * dnbinom(x, size = kmercov *
    4/bias, mu = kmercov * 4) * length)


Parameters:
         Estimate Std. Error t value Pr(>|t|)
d       1.341e-01  4.304e-03   31.16   <2e-16 ***
r       9.610e-03  8.665e-05  110.91   <2e-16 ***
kmercov 5.679e+01  7.234e-02  785.00   <2e-16 ***
bias    2.568e+00  3.679e-02   69.80   <2e-16 ***
length  2.877e+07  1.474e+05  195.15   <2e-16 ***
---
Signif. codes:  0 '***' 0.001 '**' 0.01 '*' 0.05 '.' 0.1 ' ' 1


Residual standard error: 11150 on 966 degrees of freedom


Number of iterations to convergence: 8
Achieved convergence tolerance: 6.17e-06
```
