## Supplementary figures and images for "Transcriptional and genomic parallels between the monoxenous parasite *Herpetomonas muscarum* and *Leishmania*"

### Fig S2

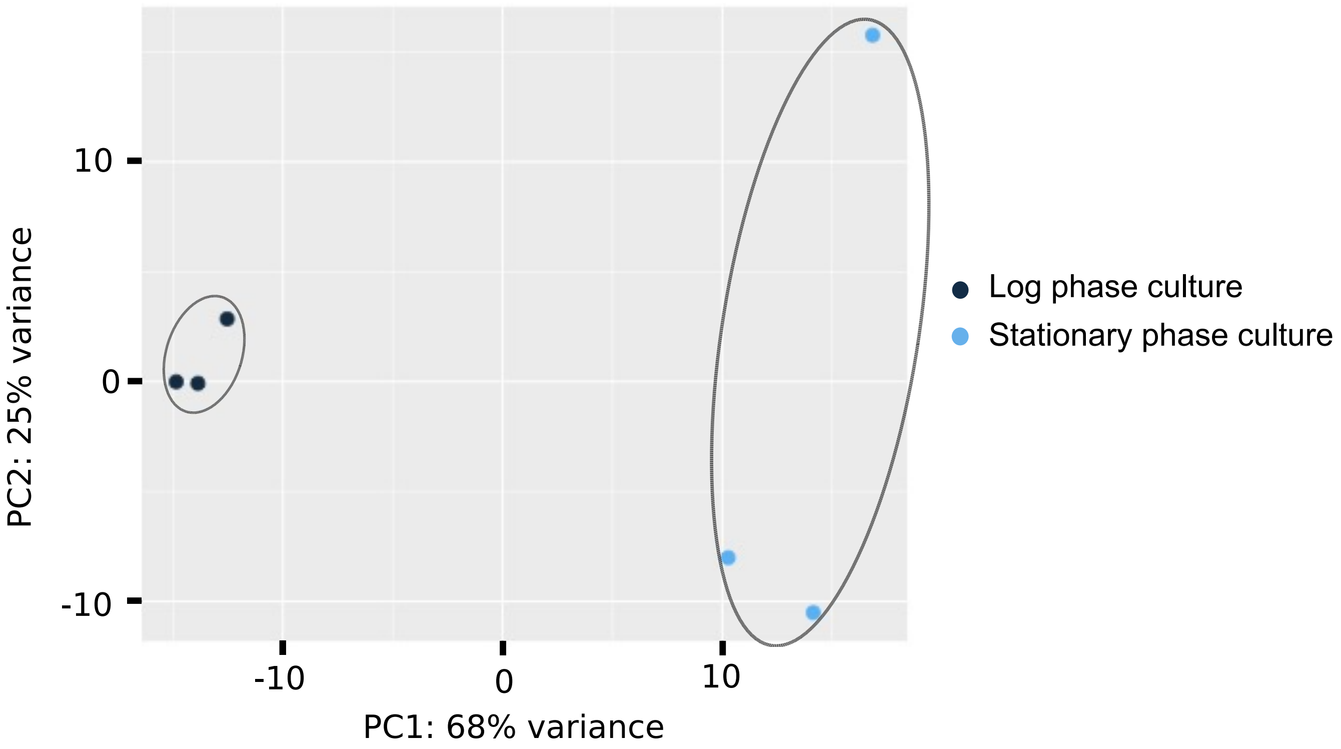
